# A machine learning method for subgroup analysis of randomized controlled trials

**DOI:** 10.1101/338996

**Authors:** Ljubomir Buturović

## Abstract

We developed a machine learning method for subgroup analyses of randomized controlled trials (RCT), and applied it to the results of the SPRINT RCT for treatment of hypertension. To date, the subgroup analyses mostly focused on detecting associations between certain factors and outcome, in the hope that the results will point out biologically (for example, carriers of a certain mutation) or clinically (for example, smokers) distinct subgroups with different outcomes. This seldom worked in the sense of re-launching the intervention for the detected subgroup only and successfully treating it. In contrast, we propose an empirical and general method to develop a predictive multivariate classifier using the RCT outcomes and baseline data. The classifier identifies patients likely to benefit from the intervention, is not limited to a single factor of interest, and is ready for validation in a subsequent pivotal trial. We believe this approach has a better chance of succeeding in identifying the relevant subgroups because of increased accuracy made possible by the use of multiple predictor variables, and opportunity to use advanced machine learning. The method effectiveness is demonstrated by the analysis of the SPRINT trial.

## Introduction

Published guidelines for subgroup analyses of randomized clinical trials favor prespecified subgroups [1,2]. The post-hoc analyses, defined as those based on examination of the RCT data, are discouraged [2]: “Such analyses are of particular concern because it is often unclear how many were undertaken and whether some were motivated by inspection of the data.” We believe that this concern unnecessarily limits the scope of what can be learned from clinical trials, and discourages discovery of subpopulations not known at the time the trial started. It also favors the hypotheses based on current knowledge, and limits opportunities for empirical learning from data. Our goal in this paper is to undertake the analyses motivated by inspection of the data. We believe that they can discover predictors of response that could be potentially beneficial to future patients, as long as the analyses are statistically sound, and findings confirmed in another clinical trial. The criticism of post-hoc analyses may also partly be due to lack of published rigorous methodology for the empirical approach. This paper aims to help filling the gap.

We propose a novel and potentially widely applicable machine learning methodology for subgroup analysis of randomized controlled trials. Our focus is on medicine, although the concept may have applications in other domains. The approach is based on developing a multivariate subgroup classifier which identifies patients likely to benefit from the given intervention. More commonly, analyses of RCTs have been based on biologically inspired features. For example, the COURAGE trial [3] showed that patients with stable coronary artery disease did not benefit from percutaneous coronary intervention (PCI). Subsequent subgroup analyses focused on multiple groups of patients identified from first or clinical principles [4–7], and found no significant benefit of the PCI treatment in any of the subgroups. Another attempt, in oncology, was more successful [8] but nevertheless also limited to confirmation of existing clinical hypothesis.

In contrast, our approach is based on empirical machine learning from data. We believe the two types of RCT analyses are significantly different, and both have merit. They correspond to distinction between empirical learning and learning from first principles, or between explanation and prediction [9], or between causal inference and correlation [10]. In this work, the focus is exclusively on the prediction (correlation) because it is worthwhile in its own right. Specifically, our goal is to develop classifier which can be applied to future patients to estimate probability of treatment benefit. Such classifier could then be validated in a subsequent prospective trial for which only patients predicted to benefit would be eligible. That way, patients could be helped even before causal understanding of the underlying process is complete. We leave methods for the causal analysis or understanding of treatment effects and study failures to separate investigations.

Previously, Simon [11] developed a PACT paradigm, focused on the RCT *design* such that the trial of a drug also produces a classifier predicting which patients benefit. This is in contrast to our work which is focused on the analysis of RCTs which already took place, and weren’t designed to yield machine learning classifiers. The latter is at present a more common situation. Despite this conceptual difference, PACT method can also be used for retrospective analysis, and an R package has been developed for that purpose [12]. Therefore we compared it with our method.

To demonstrate an implementation of our approach and evaluate its effectiveness, we applied it to the results of the SPRINT [13] randomized clinical trial for the treatment of hypertension. SPRINT showed that patients with hypertension and increased cardiovascular risk benefit from intensive treatment. The intensive treatment is defined as intervention which attempts to achieve systolic blood-pressure under 120 mm Hg. Despite the overall success, 5.2% of the SPRINT patients in the intensive treatment arm suffered cardiovascular events. This means that the intensive treatment did not benefit all individuals, and should potentially be withheld from those unlikely to benefit. Our goal was to develop a binary classifier which accepts the SPRINT trial baseline variables (demographic, clinical and laboratory variables) as input, and predicts which patients may benefit from the intensive treatment.

## Materials and methods

### Materials

The data used in analyses were based on the SPRINT clinical trial and provided through the subsequent SPRINT Data Analysis Challenge [14], organized by New England Journal of Medicine and completed in February 2017. SPRINT compared two different treatments for elevated blood pressure: intensive (systolic blood-pressure target of less than 120 mm Hg) and standard (systolic blood-pressure target of less than 140 mm Hg). We used the SPRINT baseline and outcomes data and machine learning approach to develop our algorithm. The baseline data comprised the measurements listed in Table 1, while the outcomes data comprised various endpoints, including adverse events, recorded in the trial. We used the primary endpoint to develop the machine learning classifier, and serious adverse events (SAE) endpoints for secondary analyses. The primary endpoint was a composite outcome of myocardial infarction, other acute coronary syndromes, stroke, heart failure, or death from cardiovascular causes [13]. The patients were randomly assigned by the trial to intensive and standard treatment arms.

**Table 1.** Baseline variables used to train machine learning classifiers.

| Variable | Type | Description |
| --- | --- | --- |
| Age | numeric |  |
| Aspirin | binary | Daily aspirin use |
| Female | binary |  |
| Smoking | categorical | Never, former, current |
| FRS | numeric | Framingham risk score |
| Race | categorical | Black, hispanic, white, other |
| SBP | numeric | Seated systolic blood pressure |
| DBP | numeric | Seated diastolic blood pressure |
| Agents | integer | Number of prior anti-hypertension agents |
| EGFR | numeric | Estimated glomerular filtration rate |
| SCREAT | numeric | Serum creatinine lab result |
| Cholesterol | numeric | Total cholesterol lab result |
| Glucose | numeric | Lab result |
| HDL | numeric | HDL-cholesterol lab result |
| Triglycerides | numeric | Lab result |
| UMALCR | numeric | Urine albumin lab result |
| BMI | numeric | Body mass index |
| STATIN | binary | Prior statin use |
| CVD | binary | History of clinical CVD |
| SUB | binary | History of subclinical CVD |

HDL = high-density lipoprotein, CVD = cardiovascular disease.

The race and smoking variables were one-hot encoded. All variables were standardized to zero-mean and unit standard deviation prior to machine learning.

### Methods

We sought to develop a binary classifier which assigns patients to a Positive or Negative class, using the baseline variables as inputs, such that the patients predicted Positive had, on average, large treatment benefit and those predicted Negative had on average small (or negative) benefit. The patients labeled Negative could therefore be spared the intensive treatment and its associated adverse events. This approach led to a formulation of the problem as a machine learning classification task.

The machine learning consisted of two phases: development of multiple candidate *prognostic* classifiers, and selection, among prognostic classifiers, of the best *predictive* classifier. Prognostic classifiers predict outcome of disease for a given treatment (or lack thereof), whereas predictive classifiers predict treatment benefit [15]. The core assumption of our approach was the idea that among accurate prognostic classifiers, some may also detect treatment benefit. This concept has previously been validated. For example, Oncotype DX breast cancer recurrence test was originally developed as a prognostic test [16], and subsequently shown to be predictive [17]. The idea was further supported by our own experience analyzing results of Phase III trial of Iniparib (Pathwork Diagnostics Inc. and Sanofi, unpublished data).

The first (prognostic) phase consisted in training multiple classifiers which assign probability of belonging to one of two classes: patients with event, and those without. The second phase consisted of selecting, among the prognostic classifiers, the one which optimized suitably defined clinical criterion of treatment benefit. The selection included introduction of a decision threshold which converted the probabilistic classifier into a discrete one, classifying patients as Positive or Negative. The classification enabled calculation of the clinical benefit, and therefore ranking of prognostic classifiers by the predictive metric.

Prior to machine learning, we created the training and test sets, and assigned the “true” class labels to the training samples. The training set *R* was a subset of the intensive arm. One training class contained the patients who had a primary outcome event prior to a time *T*, and the other those known not to have had an event prior to *T*. We chose *T* as the median time of all patients in the study, equal to *T* = 3.26 years, though different values could be used. The patients censored prior to T were excluded from training because we don’t know if they had an event, and thus cannot have a binary class label assigned to them.

Test set *X* was used to select the best classifier among several candidates, by clinical criteria described below. Test set was never used for any machine learning (i.e., the classification algorithms only operated on the training set). Since the selection criteria used survival analysis statistics (hazard ratios, HR), only the time of event or censoring and censorship status were needed for samples in the test set. Therefore we assigned to the test set the intensive arm patients excluded from the training set (i.e., censored), and all standard arm patients.

The resulting training and test set cardinalities were *N* = 2335 (26.7%) and *M* = 6411 (73.3%), respectively. Patients with missing data were excluded from all analyses (training and test).

To solve the machine learning problem, we trained the following classifier types (classification algorithms):

1. ElasticNet [18] binomial classifier
2. a classifier based on ridge-regularized Cox proportional hazards model [19]
3. RandomForest
4. linear Support Vector Machine (SVM)
5. PACT

The approach requires that classifiers return posterior class-conditional probability estimates which can be converted to discrete predicted labels using a decision threshold. This requirement is satisifed for state-of-the-art modern classifiers. Our methodology, described below, was applied to all algorithms except PACT. For PACT, we followed the authors’ recommendations [12].

A core question of our approach was the definition of the classifier selection statistic, i.e., the measure used to select the best classifier. The choice was based on the goal of identifying groups which differ as much as possible in terms of their treatment benefit. First step was deciding to measure the treatment benefit for a group of patients (for example, treatment benefit for all patients predicted Positive). We chose hazard ratio between intensive and standard arms, which is a natural measure for survival studies such as SPRINT (the lower the hazard ratio, the fewer events among patients assigned to the intensive vs. those assigned to the standard arms, and therefore the greater the benefit of the intensive treatment). Given this choice, a natural model selection criterion was a difference between intensive/standard hazard ratios (i.e., difference between treatment benefits) for patients predicted Positive and patients predicted Negative. Therefore, in principle we aimed to select the model which maximized the difference between intensive/standard hazard ratios for patients predicted Positive and Negative by the classifier.

However, unrestricted use of this criterion could lead to a classifier which assigns almost all patients to single class, which would have a limited clinical utility. Therefore, we imposed additional rule that a single predicted class may comprise at most a certain maximum percentage P of the overall population. In the analysis of SPRINT data, we set P = 90%, on the grounds that the clinical utility of a test which assigned over 90% of patients to either category would be low. This is a crucial element to generate a practically viable predictive classifier.

Putting these considerations together, the classifier selection rule was:

> Among classifiers which assign at most 90% of patients to one class, select the one which maximizes difference in intensive/standard arm hazard ratios between the patients predicted Positive and patients predicted Negative.

While the rule can be automatically applied, we also found it informative to visualize the data using a dotplot of the two hazard ratios, and identify best model by inspection. To avoid clutter, we only plotted the classifiers for which hazard ratio for patients predicted Positive was < 1 and the hazard ratio for patients predicted Negative was > 1. The classifiers which fail this criterion are not clinically useful.

Next we performed machine learning using the training and test data. The classifiers were trained to correctly predict the training set class using the baseline covariates as predictor variables. The training used different hyperparameter values or tuples as follows:

1. ElasticNet: (*α, λ*)
2. Random Forest: number of trees
3. linear SVM: cost
4. regularized Cox classifier: λ (cost)

The different values/tuples for each algorithm formed a one-dimensional or two-dimensional hyperparameter grid. The number of grid points for a given classifier was labeled *g_t_*, where *t* indexed the classifier type. For each algorithm, we applied exhaustive grid search by computing classifier performance at each grid point. The exhaustive search was feasible because the grids were one- or two-dimensional, with a maximum number of *g*_1_ = 200 grid points for ElasticNet. In general, other advanced classifiers such as neural networks use larger number of hyperparameters, and the computational complexity of the exhaustive grid search would be prohibitive. In those cases, random search [20] or other approaches could be used.

To convert probabilities to predictions, we applied different decision thresholds. The threshold could therefore be considered another hyperparameter. We excluded it from the grid search considerations above because the thresholding operation is very fast.

To improve classifier performance, feature selection is often applied as a step preceding classification [21]. In our initial analyses of the SPRINT data, feature selection did not provide noticeable performance gains, perhaps because of relatively few features and relatively large number of samples. Therefore it was not pursued for present research, though it remains an option for other datasets and application domains. If feature selection is used, the number of selected features becomes another hyperparameter in the grid.

To estimate classifier performance at each grid point, we used *r* = 10 repeats of 5-fold cross-validation. The purpose of the repeats was to reduce variance due to random cross-validation partition of the training set [21]. For each fold, we trained the classifier on 80% of the training data and applied it on the remaining 20%. All predictions (probabilities) from the training set, for the given repeat, were pooled together. Then, we trained the classifier on entire training set and applied it on the test set. The pooled training set probabilities were combined (pooled) with the test set probabilities and a decision threshold *D* was applied to the combined set. All samples whose probabilities exceeded the threshold were labeled (i.e., predicted) Positive, and the remaining samples were labeled Negative. We used a total of 101 thresholds, corresponding to threshold values between 0 and 1, with step 0.01. This procedure generated *S_t_* = 101 × *L_t_* sets of predicted class labels, each of cardinality *N* + *M*. This process was repeated 10 times, once for each repeat, and therefore the total number of the sets of predicted class labels equaled 10 × *S_t_* for each classifier type *t*.

As explained above, the classifier performance statistics were estimated on the pooled cross-validation predictions of the training set, and entire test set. In principle, to minimize bias one could have used just test set for this task. However, the pooling has two advantages:

1. it increases the effective number of samples used for classifier assessment
2. the test set contains a significant number of censored samples. Since we don’t know the censoring mechanism, relying exclusively on the test set may introduce confounding if the censoring is correlated with outcome. Combining the two sets reduces the impact of this because the relative contribution of censoring is reduced

In the second (predictive) phase, we computed the classifier statistics for each repeat using the predicted labels. The classifier statistics consisted of the 10 × *S_t_* triplets (*h_P_, h_N_,p*), where:

- *h_P_* = hazard ratio between intensive and standard arms for patients predicted Positive
- *h_N_* = hazard ratio between intensive and standard arms for patients predicted Negative
- *p* = proportion of patients predicted Positive

For each *S_t_*, the 10 triplets were averaged to produce mean values 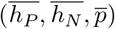. The *S_t_* triplets were plotted in a classifier dotplot. The best classifier for the type *t* was selected by applying the selection rule. Finally, the best classifiers for all types were compared to select the overall winner. The selected classifier was named *SafeSPRINT*.

The principal assessment of the SafeSPRINT performance was based on tabular presentation of key statistics (HR ratios, percent predicted Positive, and numbers of events in each arm), and Kaplan-Meier (KM) survival graphs for four groups of patients:

- Positive patients in intensive arm
- Positive patients in standard arm
- Negative patients in intensive arm
- Negative patiens in standard arm

The KM graphs required a predicted label (Positive or Negative) for each patient in the training and test sets. However, the classifier selection procedure only identified a triplet of mean values 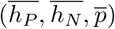 for the best classifier, not a specific set of predictions. The mean values were averaged over 10 repeats, i.e., 10 sets of predictions. To plot the KM plots, we needed to derive a single, *representative* set of predictions out of the 10 used to select the best classifier. The representative set of predictions (i.e., representative cross-validation repeat) was defined as the set of predictions corresponding to the repeat whose hazard ratios deviated least from the mean hazard ratios 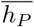 and 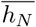. Those predictions were then used to plot the survival graphs.

The KM graphs and associated survival statistics were used to assess the expected performance of the classifier in a future, independent validation. For a successful classifier, the expectation was that the Positive patients should exhibit better event-free survival in the intensive arm, whereas Negative patients should exhibit better event-free survival in the standard arm.

The pseudo-code for the method to find the best classifier for identifying the population of subjects benefiting from a treatment is shown in Algorithm 1 box. The functions *f*(), *x*() and *z*() referenced therein are well-known machine learning [22] and survival analysis [23] algorithms. After the algorithm returns best hyperparameters *b*, it is straightforward to train the final classifier using *R, M* and *H_b_*.

**Algorithm 1.**
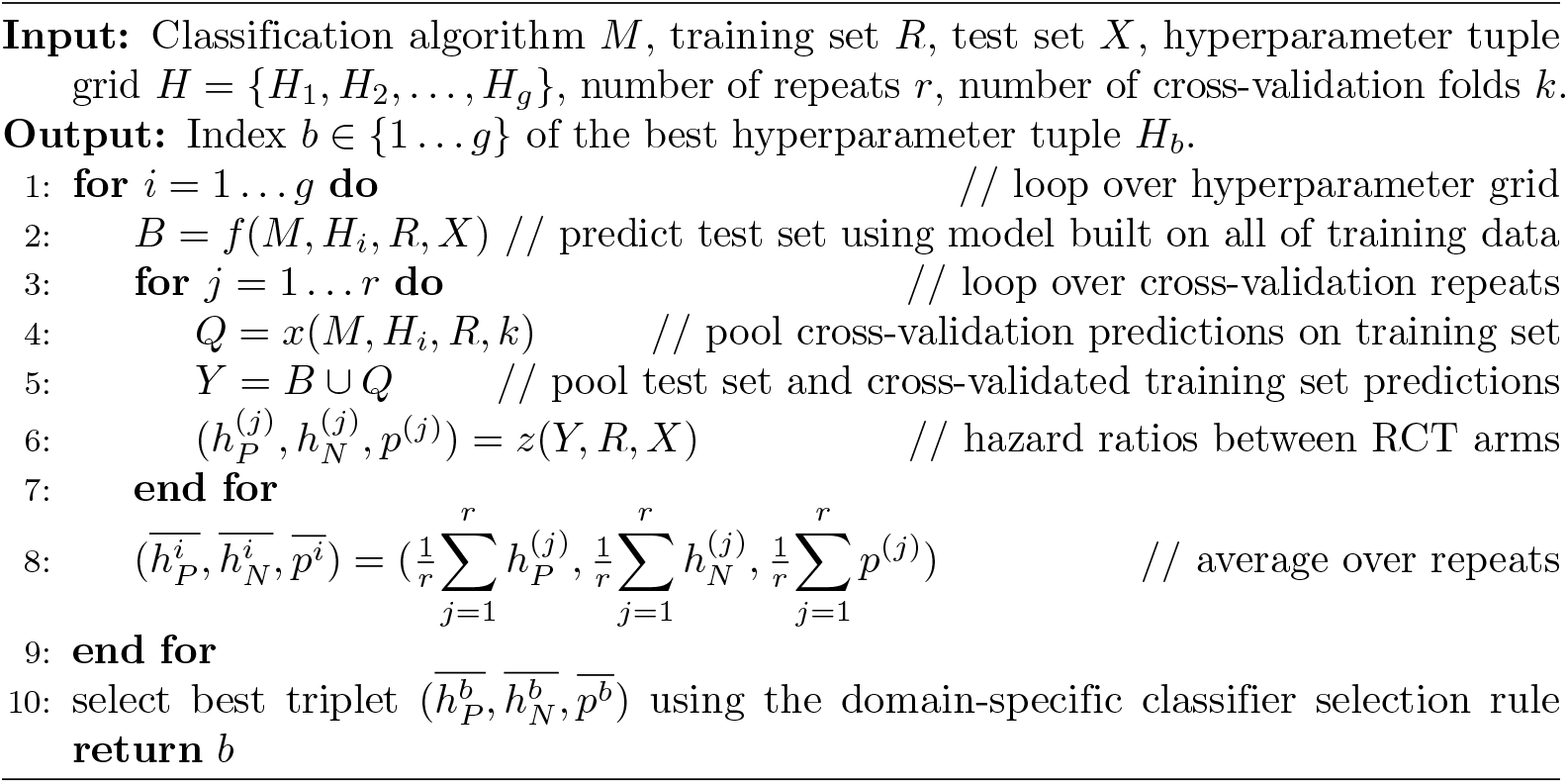
Develop a classifier for identifying the population of subjects benefiting from a treatment in an RCT. *R* and *X* are matrices of features, class label, RCT arm indicator and survival data. *H* is hyperparameter grid for classification algorithm *M*, plus the decision threshold. *f*() trains *M* on *R* using *H*, and returns predicted labels for samples in *X*. *x*() performs random *k*-fold cross-validation using *M, R* and *H*, and returns pooled cross-validated predicted labels *Y* for samples in *R*. *z*() estimates hazard ratios between the RCT arms for Positive and Negative samples in *Y*, using arm labels and survival data from *R, X. p* is proportion of samples predicted Positive.

Per literature recommendations [1,2], we also performed likelihood ratio test of interaction between label assigned by SafeSPRINT, and treatment arm assignment, and generated the related forest plot. We did not apply multiple-comparison adjustment because the interaction test was performed exactly once during this research project (i.e., there were no multiple comparisons).

We paid particular attention to minimizing overfitting and avoiding false positive conclusions in our analyses. To that end, each training set sample was predicted in cross-validation mode, meaning that it was not used in the learning of the cross-validation model which predicted its outcome. Predictions were performed 10 times, to minimize bias resulting from different cross-validation partitionings. The test set samples, representing 73.3% of all data, were used exclusively to select the best model and never in any learning. The Kaplan-Meier assessment of the classifier was only performed once after it was locked. Finally, the p-value of the interaction between SafeSPRINT risk assignment and treatment arm was computed exactly once, after the classifier was locked and all sample predictions produced.

For PACT, we followed the software recommendations and generated the KM graphs for patients predicted to benefit vs. those predicted not to benefit. The results were compared with SafeSPRINT.

### ElasticNet

The winning classifier was an ElasticNet classifier (see “Results”). The ElasticNet hyperparameters are *α*, which governs the trade-off between the lasso and ridge regularizations, and *λ*, which controls the magnitude of the weight regularization. We ran ElasticNet with *α* = 0, which corresponds to the ridge classifier, and *α* = 1, which corresponds to the lasso classifier. The lasso classifier had a poor binary classification performance, and was removed from further analyses. A set of *g*_1_ *λ* values for the ridge classifier was determined using the standard procedure defined by the ElasticNet implementation “glmnet” [18]. We set *g*_1_ (the number of *λ* values) to 100, which is the default value in “glmnet”. Each classifier was trained using the ElasticNet optimization algorithm until convergence.

### Results

We applied the proposed analysis method to the SPRINT data. To illustrate the classifier selection rule and highlight large differences among classifier types, we generated classifier dotplot graphs for ElasticNet, Cox and SVM. No graph was generated for Random Forest because no Random Forest classifier met our requirement that the Positive hazard ratio must be < 1 and Negative hazard ratio must be > 1. The graphs are shown in Figs 1, 2 and 3.

**Fig 1.**
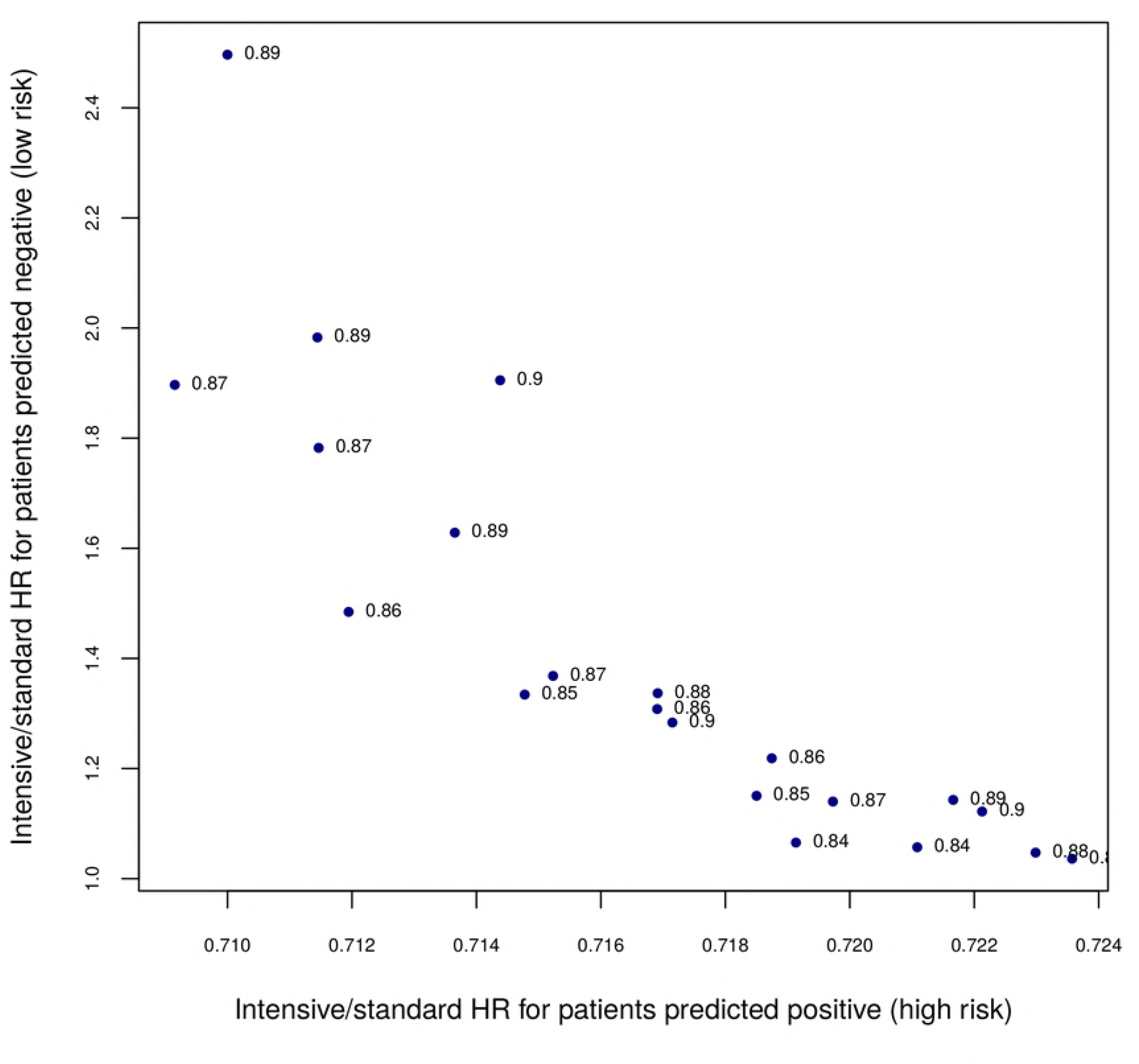
Classifier graph for ElasticNet. Each point represents a hyperparameter tuple. Only qualifying classifiers (tuples) are shown. The coordinates are average values of 10 cross-validation repeats. The numbers next to the points are proportion predicted Positive, averaged over the 10 repeats. The best classifier corresponds to the point in the upper left corner. See text for details.

**Fig 2.**
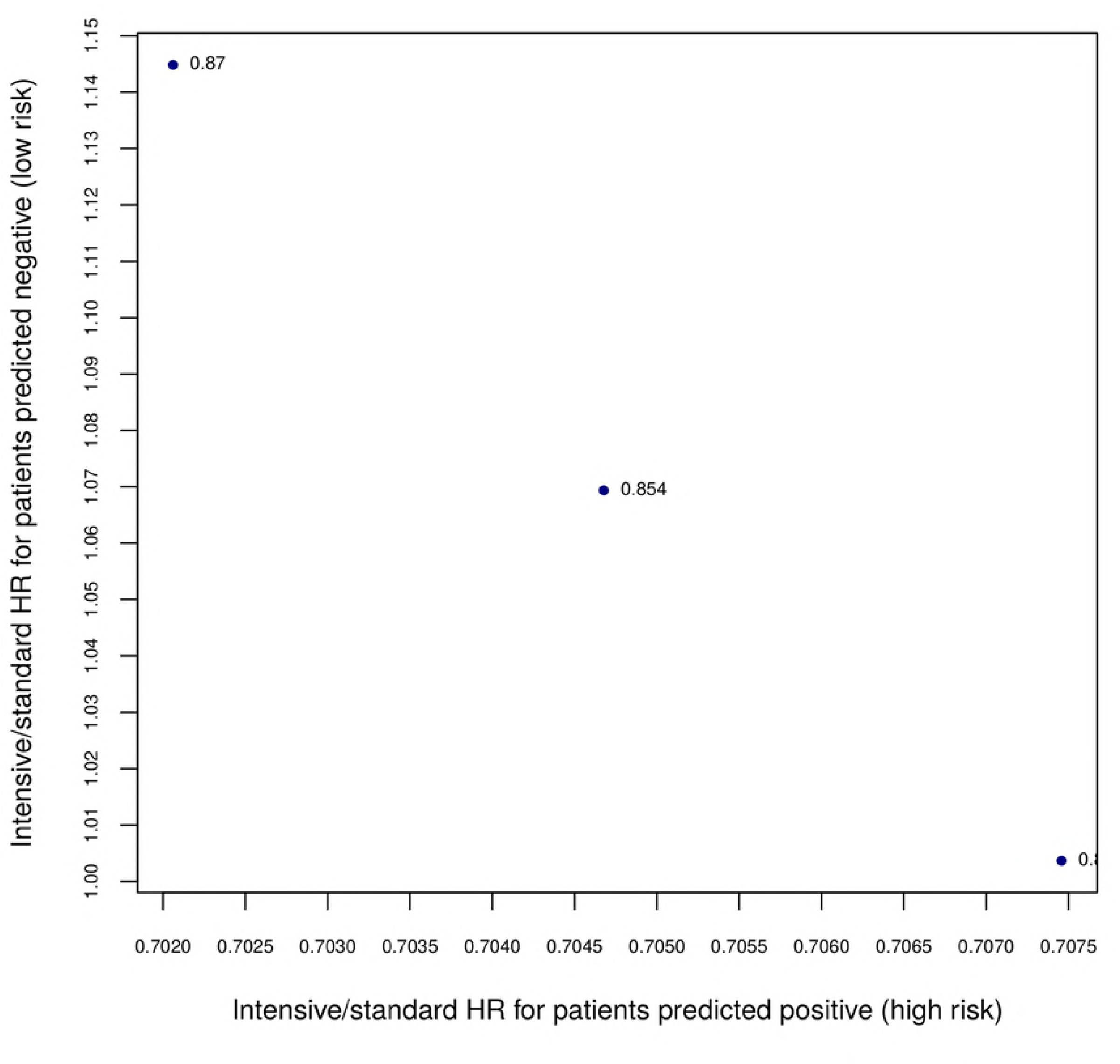
Classifier graph for Cox-based classifier.

**Fig 3.**
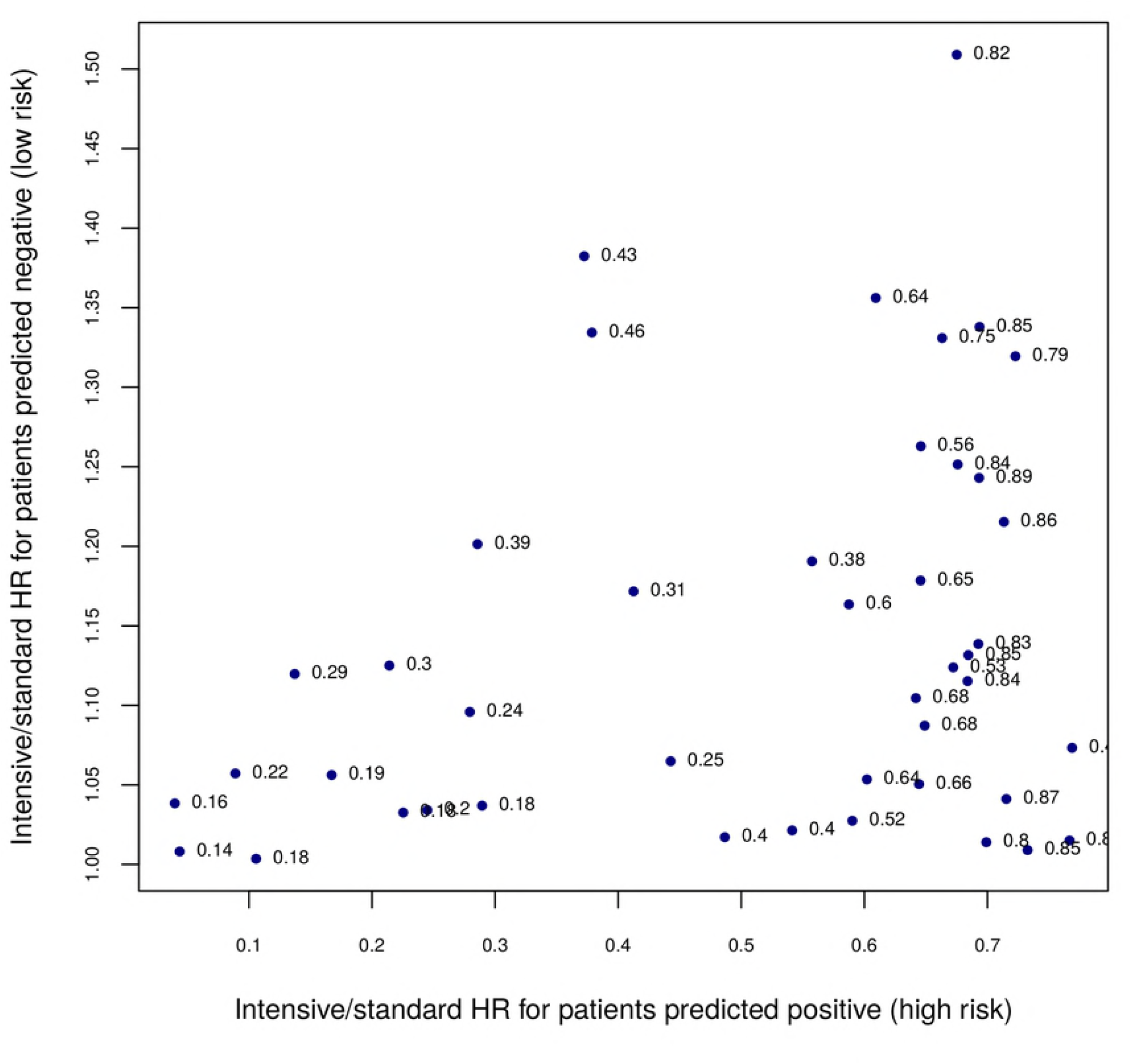
Classifier graph for SVM classifier.

The PACT results are shown in Fig4. It suggests that the classifier was not able to separate a subgroup of patients who did not benefit from treatment from those who did. In fact, the group predicted not to benefit, benefited more.

**Fig 4.**
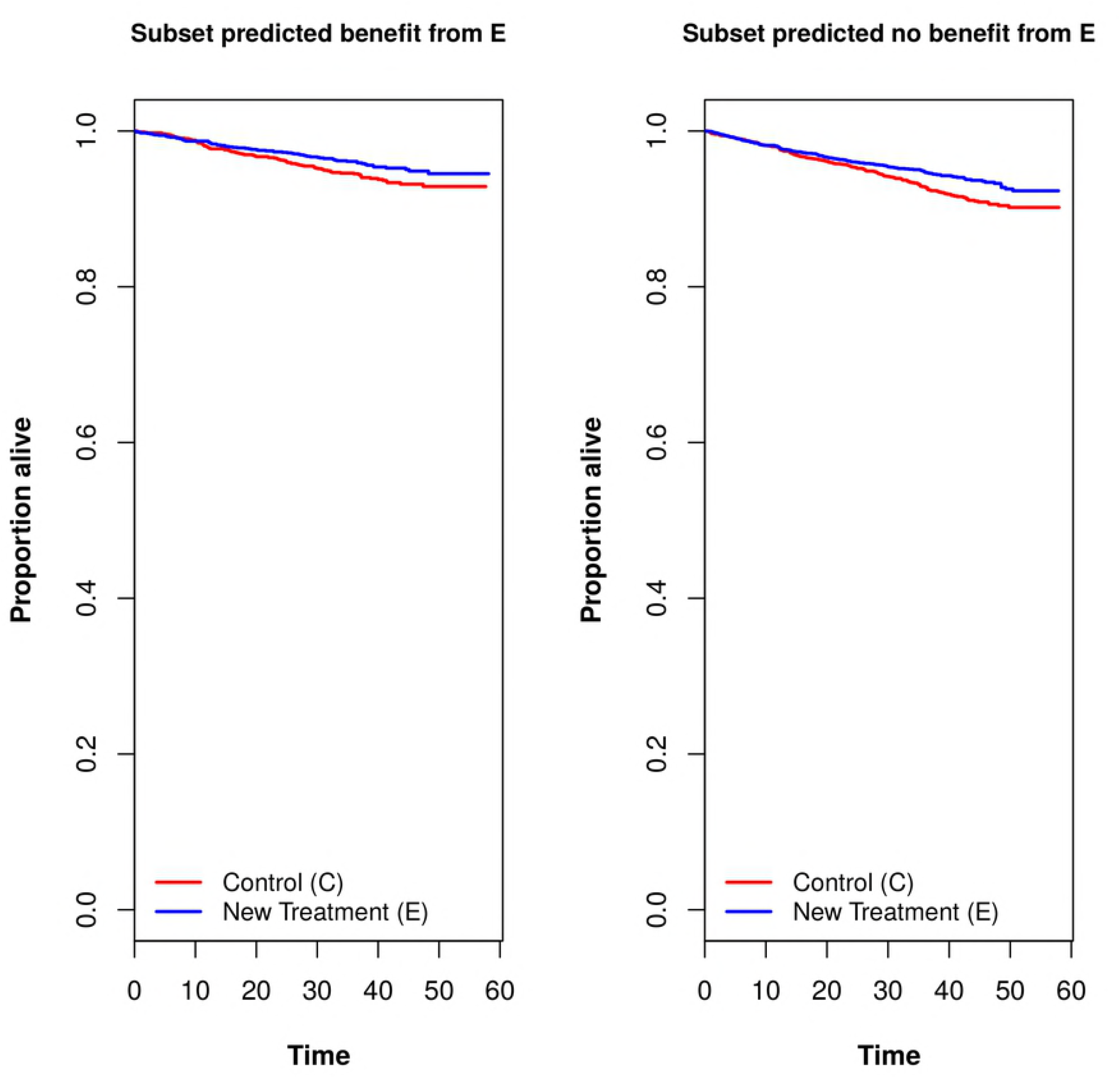
Kaplan-Meier graphs for a classifier based on PACT method. E = intensive arm; C = standard arm.

By visual inspection of Figs 1, 2, 3 and 4, it was clear that ElasticNet classifier type (Fig 1) was best by a wide margin. Among the ElasticNet classifiers, the best by our criteria corresponded to the upper leftmost point in Fig 1, because it had biggest HR gap between patients predicted Positive and patients predicted Negative.

The best classifier (SafeSPRINT) assigned 89% of the patients to Positive category and 11% to Negative. The key statistics of SafeSPRINT are shown in Table 2.

**Table 2.**
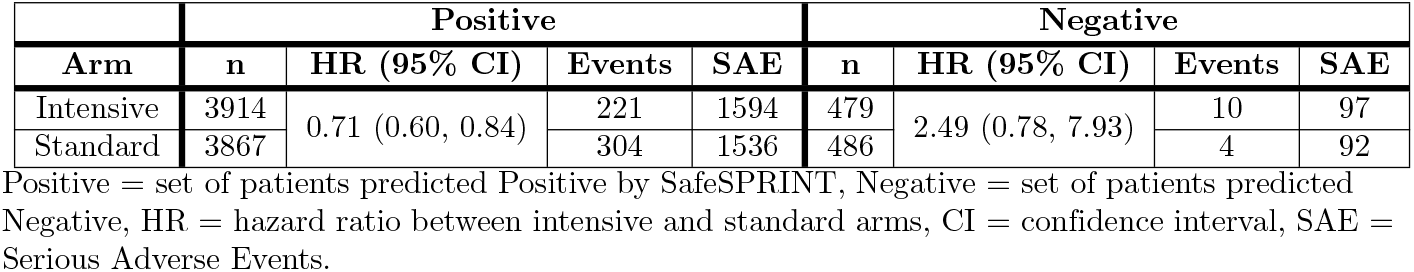
Key statistics of the SafeSPRINT classifier.

| Arm | Positive |  |  |  | Negative |  |  |  |
| --- | --- | --- | --- | --- | --- | --- | --- | --- |
|  | n | HR (95% CI) | Events | SAE | n | HR (95% CI) | Events | SAE |
| Intensive | 3914 | 0.71 (0.60, 0.84) | 221 | 1594 | 479 | 2.49 (0.78, 7.93) | 10 | 97 |
| Standard | 3867 |  | 304 | 1536 | 486 |  | 4 | 92 |

Positive = set of patients predicted Positive by SafeSPRINT, Negative = set of patients predicted Negative, HR = hazard ratio between intensive and standard arms, CI = confidence interval, SAE = Serious Adverse Events.

Table 2 shows the key result: patients predicted Positive had more events in the standard arm, whereas patients predicted Negative had more events in the intensive arm.

The relative frequency of SAE among Negative patients (19.6%) was approximately half that among Positive patients (40.2%). The rates for both subgroups were approximately the same in both treatment arms.

To further assess benefit, or lack thereof, of intensive hypertension treatments for the two predicted groups of patients, we plotted Kaplan-Meier estimates of probabilities of primary outcomes (survival functions) in four groups of patients: predicted Positive in the intensive and standard arms, and predicted Negative in the intensive and standard arms. Fig 5 shows the survival curves for the four groups.

**Fig 5.**
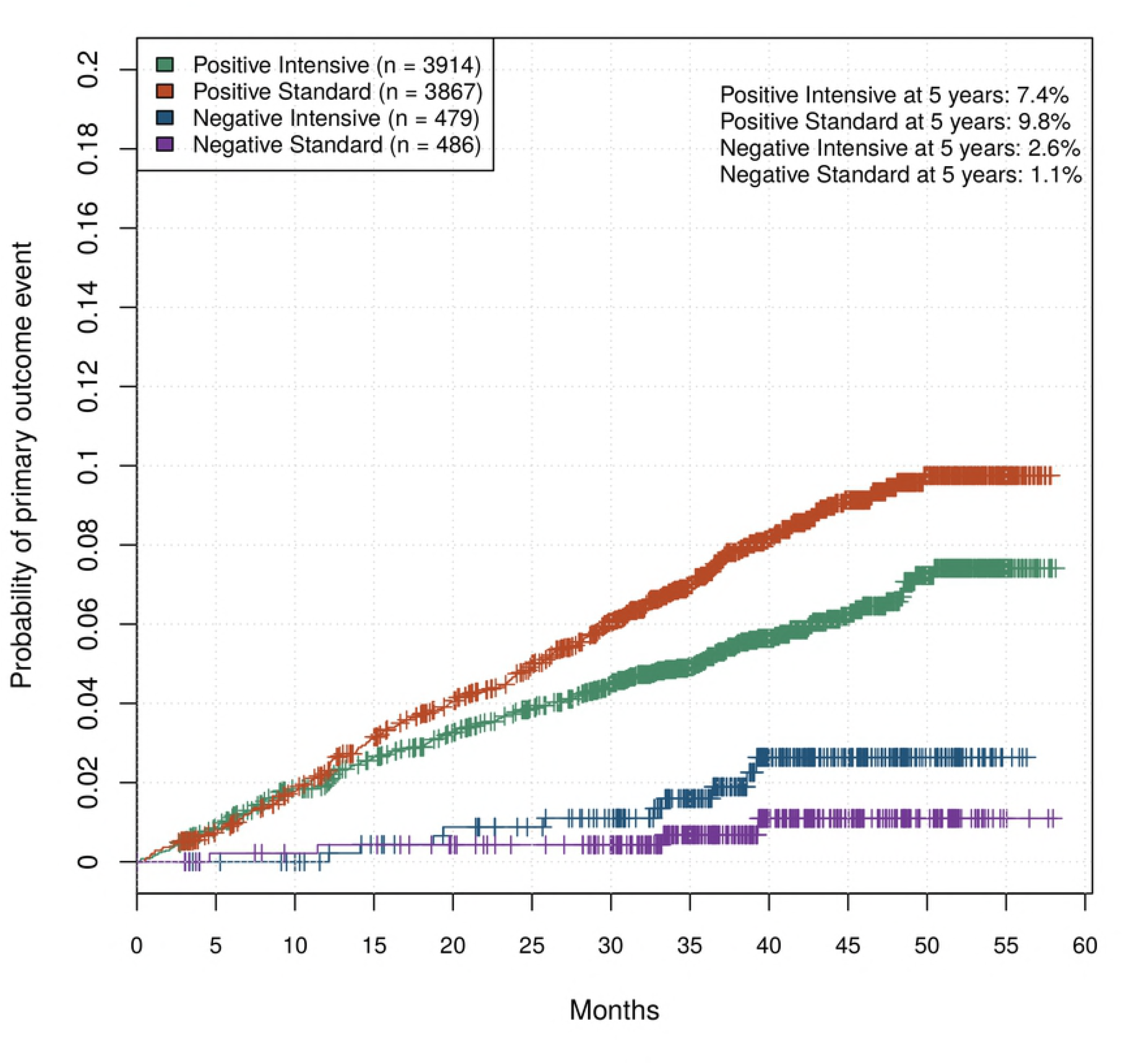
Probability of events for Positive and Negative patients under the standard and intensive treatments.

Fig 5 visualizes and amplifies the trends already observed in Table 2: the intensive treatment reduces number of events in the Positive group, and increases number of events in the Negative group. In relative terms, the nominal effect is much stronger in the Negative group because the number of events and the probability of event at 5 years more than doubled with intensive care. However, it is not statistically significant due at least in part to smaller size of the Negative group.

We performed the likelihood ratio test of interaction between SafeSPRINT category and treatment arm. The null hypothesis is that the efficacy of the treatment is the same among Positive and Negative patients. The P value was 0.02. The corresponding forest plot is shown in Fig 6. It suggests that the interaction is potentially qualitative [1] (i.e., that the treatment is beneficial for one subgroup and harmful for the other), which is of particular clinical interest.

**Fig 6.**
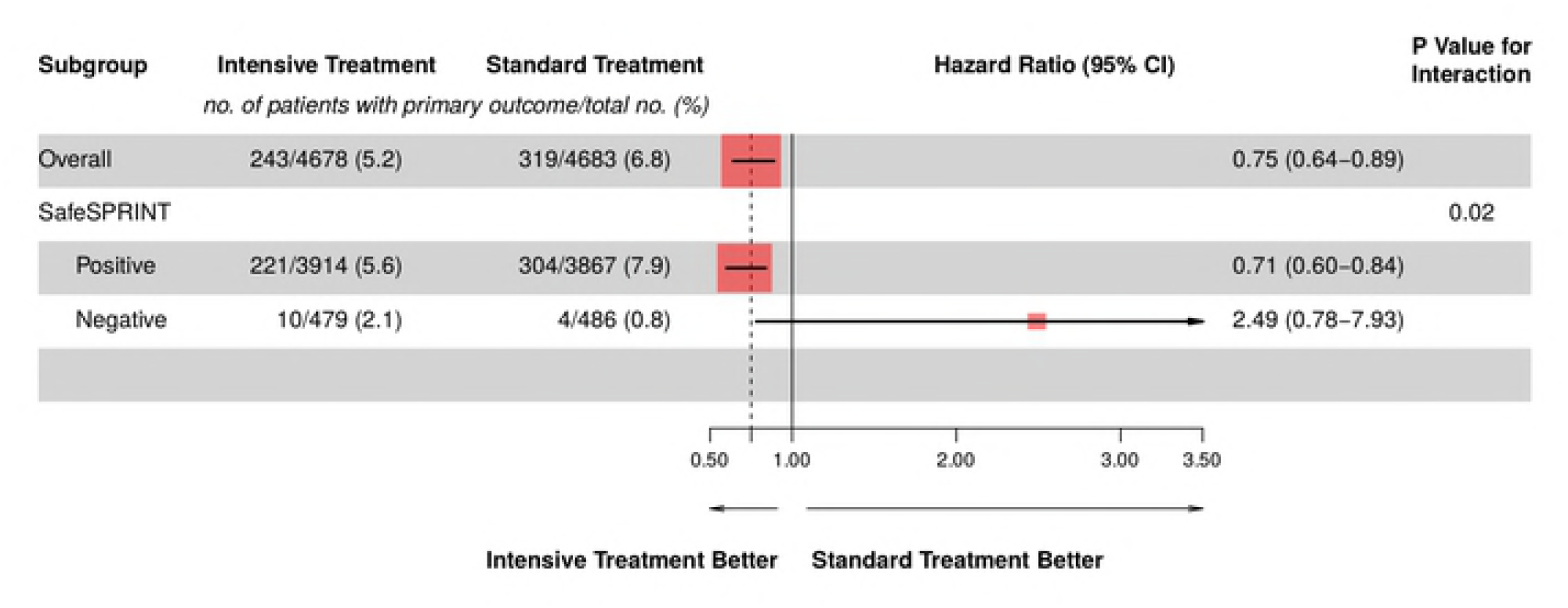
Effect of SPRINT treatment on Positive and Negative patients as defined by our classifier. The size of the colored squares is proportional to number of samples in the corresponding group.

## Discussion

SafeSPRINT-positive patients benefited from the Intensive treatment, as expected, since they represent almost 90% of the SPRINT trial population, and the population as a whole did benefit. The benefit was statistically significant. In contrast, Negative patients had more primary events in the Intensive arm than in Standard, but the effect was not statistically significant. This could be because SPRINT was not powered to detect such subgroups, or because the effect in fact wasn’t significant, or both. Considering the magnitude of the difference, and the fact that the interaction test was statistically significant, and the fact that the analyses were carefully planned to minimize bias, our interpretation is that the SafeSPRINT-negative patients probably do not benefit from the intensive treatment, and could be harmed. If this is proven, they should preferably adhere to the standard systolic blood-pressure target of 140 mm Hg. However, only an adequately-powered randomized controlled trial could conclusively confirm this hypothesis.

Surprisingly, we also found that Negative patients had much lower rate of serious adverse events (19.6% vs. 40.2% for Negative and Positive patients, respectively). This is particularly interesting given that adverse events were not part of the classifier training. Based on this observation, the SafeSPRINT Negative group can be characterized as patients who are potentially harmed by the intensive treatment, but are nonetheless overall much less sensitive to other side effects of hypertension treatment. This finding suggests that SafeSPRINT could also be used as a prognostic classifier of side-effects.

An advantage of our empirical learning approach is potentially greater accuracy in predicting subgroup benefit, and consequently better patient outcomes. A limitation is increased model complexity and limited interpretability. In the event, the chosen classifier was linear (ElasticNet), which is amenable to interpretation, therefore the limitation did not materialize. In general, however, the chosen classifier might be non-linear, and thus harder to interpret.

We note that although our proposed method was partially motivated by the SPRINT Challenge, it could also be applied to analyses of other clinical trials, and potentially to other randomized control trials, outside of medicine. In medicine, analyses of failed oncology RCTs could use a very similar approach because the goal is similar: finding a subgroup of patients who maximally benefit from the new treatment.

Our method outperformed previously published algorithm PACT by a wide margin. One possible explanation is that the PACT software package supports a limited family of classifiers. Also, it is possible that it does not perform advanced hyperparameter search.

The SPRINT trial design and results have been subject to criticism [24]. Nevertheless, to date it remains the best available evidence on the merits of the specific blood-pressure lowering targets. Furthermore, the principled analysis approach proposed in this paper remains applicable regardless of whether clinical community eventually accepts the SPRINT treatment guidelines.

The principal limitation of this research is that the proposed SafeSPRINT hypothesis, and therefore the method by which it was generated, have not been independently validated. Given the cost and complexity of executing randomized controlled trials in medicine, this is a practical limitation which cannot easily be overcome. However, in other fields such as advertising, the independent validation may be feasible.

Another limitation is that our classifier selection criterion did not include the frequency of SAE. Ideally, this outcome should be made part of the selection rule such that the best classifier minimizes a composite measure of primary events and SAE. However, that would require quantifying the relative importance and impact of the SAE and primary outcomes in cardiovascular disease, using metrics such as quality of life. During the project, we did not have access to adequate clinical expertise to do this, and therefore development of machine learning classifiers which account for the adverse events are left for future research. We believe that the proposed framework is sufficiently general to include adverse events.

We used advanced core classifiers Random Forest, ElasticNet, SVM and regularized Cox, but not other state-of-the-art classifiers like XGBoost [25], non-linear SVM and neural networks. The reason is that at the time of this research we have not completed integration of these classification algorithms into our analysis procedure. This is a limitation and an opportunity for improvement of the SafeSPRINT performance. It is is left for further research.

While elements of these ideas have been put forward before, to the best of our knowledge no single reference described a generic machine learning subgroup analysis procedure, demonstrated the performance of the method on a real and significant clinical trial, and provided the tools to repeat the results.

## Conclusion

In this paper, we proposed an empirical procedure for deriving predictive hypothesis (machine learning classifier) from the results of randomized controlled trials. Such a classifier could be validated in subsequent trial(s), and then used to assign the treatment to people most likely to benefit, and withhold it from people unlikely to benefit.

The main contributions of the paper are:

- proposed the idea of separating the causal and correlative (empirical) goals in the subgroup analyses of randomized controlled trials
- using the empirical approach, defined a general and relatively easily implementable procedure for machine learning analysis of subgroups
- demonstrated the implementation and effectiveness of the proposed approach by applying it to the results of SPRINT randomized clinical trial
- formulated a data-supported hypothesis that a subgroup of 11% of patients with hypertension do not benefit from the intensive treatment, targeting systolic blood-pressure under 120 mm Hg
- showed that our method performs significantly better than the previously published work on this subject

We hope these contributions should motivate the empirical, machine-learning approach to subgroup analyses of future trials.

## Acknowledgments

The author wishes to thank Damjan Krstajić for helpful discussions and Alejandrina Pattin for software development.

## References

1. Rothwell PM. Subgroup analysis in randomised controlled trials: importance, indications, and interpretation. Lancet 2005 Jan;365:176–86.

2. Wang R, Lagakos SW, Ware JH, Hunter JD, Drazen MJ. Statistics in medicine – reporting of subgroup analyses in clinical trials. New Engl J Med. 2007 Nov;357(21):2189–2194.

3. Boden WE, O’Rourke RA, Koon KT, Hartigan PM, Maron DJ, Kostuk WJ, et al. Optimal medical therapy with or without PCI for stable coronary disease. New Engl J Med. 2007 Apr;356:1503–1516.

4. Shaw LJ, Weintraub WS, Maron DJ, Hartigan PM, Hachamovitch R, Min JK, et al. Baseline stress myocardial perfusion imaging results and outcomes in patients with stable ischemic heart disease randomized to optimal medical therapy with or without percutaneous coronary intervention. Am Heart J. 2012 Aug;164(2):243–50.

5. Mancini JGB, Bates ER, Maron DJ, Hartigan P, Dada M, Gosselin G, et al. Quantitative results of baseline angiography and percutaneous coronary intervention in the COURAGE trial. Circ Cardiovasc Qual Outcomes. 2009 Jul;2(4):320–327.

6. Spertus JA, Maron DJ, Cohen DJ, Kolm P, Hartigan P, Weintraub WS, et al. Frequency, predictors, and consequences of crossing over to revascularization within 12 months of randomization to optimal medical therapy in the clinical outcomes utilizing revascularization and aggressive drug evaluation (COURAGE) trial. Circ Cardiovasc Qual Outcomes. 2013 Jul;6(4):409–418.

7. Mancini GB, Hartigan PM, Bates ER, Chaitman BR, Sedlis SP, Maron DJ, et al. Prognostic importance of coronary anatomy and left ventricular ejection fraction despite optimal therapy: assessment of residual risk in the Clinical Outcomes Utilizing Revascularization and Aggressive DruG Evaluation Trial. Am Heart J. 2013 Sep;166(3):481–7.

8. Liu S, Bingshu C, Burugu S, Leung S, Dongxia G, Shakeel V, et al. Role of Cytotoxic Tumor-Infiltrating Lymphocytes in Predicting Outcomes in Metastatic HER2-Positive Breast Cancer: A Secondary Analysis of a Randomized Clinical Trial. JAMA Oncol. 2017 Nov. doi: 10.1001/jamaoncol.2017.2085.

9. Casti JL. Searching for certainty: what scientists can know about the future. 1st ed. New York: William Morrow and Company, Inc.; 1990.

10. Pearl J, Glymour M, Jewell NP. Causal inference in statistics: a primer. 1st ed. Chichester, United Kingdom: John Wiley & Sons Ltd; 2016.

11. Simon R. Clinical trials for predictive medicine. Statist Med. 2012 Jun;31:3031–3040.

12. Subramanian J, Simon R. pact: Predictive Analysis of Clinical Trials; 2016. Available from: https://CRAN.R-project.org/package=pact.

13. The SPRINT Research Group. A randomized trial of intensive versus standard blood-pressure control. New Engl J Med. 2015 Nov;373(22):2103–2116.

14. Burns NS, Miller PW. Learning what we didn’t know – the SPRINT data analysis challenge. New Engl J Med. 2017 Jun;376:2205–2207.

15. Simon R. Advances in clinical trial designs for predictive biomarker discovery and validation. Curr Breast Cancer Rep. 2009 Dec;1(4):216–221.

16. Paik S, Shak S, Tang G, Kim C, Baker J, Cronin M, et al. A multigene assay to predict recurrence of tamoxifen-treated, node-negative breast cancer. New Engl J Med. 2004 Dec;351(27):2817–26.

17. Albain KS, Barlow WE, Shak S, Hortobagyi GN, Livingston RB, Yeh I-T, et al. Prognostic and predictive value of the 21-gene recurrence score assay in a randomized trial of chemotherapy for postmenopausal, node-positive, estrogen receptor-positive breast cancer. Lancet Oncol. 2010 Jan;11(1):55–65.

18. Friedman J, Hastie T, Tibshirani R. Regularization paths for generalized linear models via coordinate descent. J Stat Softw. 2010 Jan;33(1):1–22.

19. Bovelstad HM, Nygard S, Storvold HL, Aldrin M, Borgan O, Frigessi A, et al. Predicting survival from microarray data – a comparative study. Bioinformatics. 2007 Aug;23(16):2080–2087.

20. Bergstra J, Bengio Y. Random search for hyper-parameter optimization. J Mach Learn Res. 2012;13:281–305.

21. Krstajic D, Buturovic LJ, Leahy DE, Thomas S. Cross-validation pitfalls when selecting and assessing regression and classification models. J Cheminform. 2014 Apr. doi: 10.1186/1758-2946-6-10.

22. Hastie T, Tibshirani R, Friedman J. The elements of statistical learning: data mining, inference and prediction. 2nd ed. New York: Springer; 2016.

23. Kleinbaum DG, Klein M. Survival analysis. 2nd ed. New York: Springer; 2005.

24. Husten L. New questions raised about SPRINT. CardioBrief. 2017. Available from: http://www.cardiobrief.org/2017/02/08/new-questions-raised-about-sprint

25. Chen T, Guestrin C. XGBoost: A scalable tree boosting system. ACM SIGKDD Conf. Knowledge Discovery and Data Mining 2016 Aug. doi: 10.1145/2939672.2939785.

